# Systematic Prediction of Regulatory Motifs from Human ChIP-Sequencing Data Based on a Deep Learning Framework

**DOI:** 10.1101/417378

**Authors:** Jinyu Yang, Adam D. Hoppe, Bingqiang Liu, Qin Ma

**Affiliations:** Bioinformatics and Mathematical Biosciences Lab, Department of Agronomy, Horticulture and Plant Science, South Dakota State University, BioSNTR, Brookings, SD, USA, 57007; Department of Mathematics and Statistics, South Dakota State University, Brookings, SD, USA, 57007; Department of Chemistry and Biochemistry, Avera Health and Sciences Building, RM 131 South Dakota State University (SDSU), BioSNTR, Brookings, SD 57007; School of Mathematics, Shandong University, Jinan, China, 250100

**Keywords:** TF-DNA interactions, ChIP-Seq, Motif prediction, Deep learning, DNA shape motif

## Abstract

Identification of transcription factor binding sites (TFBSs) and *cis*-regulatory motifs (*motifs* for short) from genomics datasets, provides a powerful view of the rules governing the interactions between TFs and DNA. Existing motif prediction methods however, are limited by high false positive rates in TFBSs identification, contributions from non-sequence-specific binding, and complex and indirect binding mechanisms. High throughput next-generation sequencing data provides unprecedented opportunities to overcome these difficulties, as it provides multiple whole-genome scale measurements of TF binding information. Uncovering this information brings new computational and modeling challenges in high-dimensional data mining and heterogeneous data integration. To improve TFBS identification and novel motifs prediction accuracy in the human genome, we developed an advanced computational technique based on deep learning (DL) and high-performance computing, named DESSO. DESSO utilizes deep neural network and binomial distribution to optimize the motif prediction. Our results showed that DESSO outperformed existing tools in predicting distinct motifs from the 690 *in vivo* ENCODE ChIP-Sequencing (ChIP-Seq) datasets for 161 human TFs in 91 cell lines. We also found that protein-protein interactions (PPIs) are prevalent among human TFs, and a total of 61 potential tethering binding were identified among the 100 TFs in the K562 cell line. To further expand DESSO’s deep-learning capabilities, we included DNA shape features and found that (*i*) shape information has a strong predictive power for TF-DNA binding specificity; and (*ii*) it aided in identification of the shape motifs recognized by human TFs which in turn contributed to the interpretation of TF-DNA binding in the absence of sequence recognition. DESSO and the analyses it enabled will continue to improve our understanding of how gene expression is controlled by TFs and the complexities of DNA binding. The source code and the predicted motifs and TFBSs from the 690 ENCODE TF ChIP-Seq datasets are freely available at the DESSO web server: http://bmbl.sdstate.edu/DESSO.

## INTRODUCTION

Gene regulation is essential for the versatility and adaptability of an organism’s cellular and biochemical pathways to meet the demands of its environment. Gene expression is regulated by a set of transcriptional regulatory signals, including TFs, microRNAs, long non-coding RNA, and epigenomic regulators [1, 2]. TFs control gene expression by binding to specific DNA sequences, i.e., TFBSs [3], and the aligned profiles of TFBSs are referred to as *cis*-regulatory motifs [4, 5]. The binding or releasing of TFs promote or suppress transcription to guarantee the target genes are expressed at the proper time and in the appropriate amount according to particular cell states and circumstance [6, 7]. Substantial efforts have been invested in the studies of TF binding specificities and motif prediction, resulting in the development of numerous algorithms and computational tools and databases [8-17]. However, the understanding of TF-DNA binding mechanism is still fragmented, and its computational elucidation is still a considerable challenge in systems biology [18-21].

Beyond the sequence level, recent studies have highlighted the essential role of DNA structure in quantitatively influencing TF-DNA binding specificity both *in vitro* and *in vivo* across diverse TF families [22-25]. Owing to the advances in DNA structure elucidation, four distinct DNA shape features (i.e., Minor Groove Width (MGW), Propeller Twist (ProT), Helix Twist (HelT), and Roll) can be computationally derived from DNA sequences by Monte Carlo simulation [26]. These features, if experimentally validated, can be considered as shape motifs and thereby an extension of traditional regulatory sequence motifs. This idea is corroborated by the fact that the flanking regions of TFBSs can contribute to the binding specificity indirectly by influencing the DNA shape of the target sequences. Thus, features in DNA sequences in combination with shapes determine the TF binding in a more sophisticated way than was originally thought [27- 30].

ChIP-Seq provides the genome-wide interactions between DNA and DNA-associated proteins and large-scale ChIP-Seq efforts enable new insights into gene regulation analyses. A considerable amount of ChIP-Seq data has been generated in the public domain, including approximately 6,000 datasets of human from the ENCODE database [31]. These datasets provide an unprecedented opportunity to predict motifs, identify TFBSs, and capture more features affecting TF binding [32]. ChIP-Seq data mining and modeling have many challenges in computation, facing high-dimensional and heterogeneous data properties, and a variety of popular methods have been developed [33-38]. However, computational challenges remain for the accurate and exhaustive identification of motifs [38].

Deep Learning (DL) has achieved unprecedented performance, among the big data methods, for capturing motif patterns and elucidating complex regulatory mechanisms [39, 40]. Specifically, DL methods enable extraction of more complex patterns than rational models owing to their multi-layer network architecture. Importantly, the intrinsic overfitting issue in multi-layer DL networks can be overcome by the large scale of the available datasets [41-44]. For example, the convolutional neural network (CNN) in DL improved the state-of-the-art performance in TF-DNA binding specificity prediction through optimizing the position weight matrix (PWM)-like motif detectors [45]; and the motif patterns predicted by DeepBind have been well mapped to documented motifs by Alipanahi *et al*. based on available human TFBS databases [41]. However, these methods focused on predicting TF-DNA binding specificity and failed to identify motifs accurately, giving rise to high false positive and false negative issues (26).

Identifying where a TF binds is very important for gene regulation-related studies, including but not limited to the occurrence and development of complex diseases and cell proliferation and differentiation. Furthermore, studies about why the TF binds to specific sites (sometimes not conserved in sequence level) are essential for elucidating the underlying mechanisms of regulation [3]. Hence, the demand for the following data analysis and interpretation is significant: (*i*) how to improve the performance of capturing sequence-specific TFBSs; and (*ii*) how the non-sequence-specific TFBSs contribute to TF-DNA binding in alternative ways. DNA shape features can be integrated into DL models to explore how these features quantitatively contribute to TF-DNA binding, even though conserved shape patterns (or shape motifs) encoded in the human genome are still not well modeled (34,35). Furthermore, in-depth analysis of novel identified motif patterns containing both sequence and shape factors, will facilitate new insight into hypothetical mechanism regulating TF binding that can be experimentally validated.

In this study, we proposed a novel DL framework, named DESSO (<u>DE</u>ep <u>S</u>equence and <u>S</u>hape m<u>O</u>tif), using the CNN model to predict motifs and identify TFBSs in both base pair and regional DNA shape features. The identified motifs are evaluated using documented motifs in JASPAR [16] and TRANSFAC [15], compared with DeepBind [41], Basset [43], MEME-ChIP [35], and gkm-SVM [46]. Further analyses were conducted by integrating multiple biological information including TF binding domain types, chromatin accessibility, phylogenetic conservation, PPI, *etc*. For the first time, rather than determine motif instances (i.e., TFBSs) using a subjective cutoff as previously reported [41-43], we integrated the binomial distribution into DESSO to optimize the motif instances identification based on identified motif patterns [47]. DESSO was evaluated using 690 *in vivo* ENCODE ChIP-Seq datasets [48], and the results demonstrated that the motifs generated by DESSO captured the binding specificity of TFs more accurately than the other four tools. Meanwhile, 61 indirect binding activities were identified based on these motifs, some of which were confirmed as known tethering binding activities and PPI. The combination of four DNA shape features was used in DESSO to investigate their contribution in predicting TF-DNA binding specificity. The shape information provided an additional dimension to explain TF binding even without sequence specificity, and the identified shape motifs uncovered that functionality conserved shape patterns are common in the human genome.

## RESULTS

### DESSO (DEep Sequence and Shape mOtif) pipeline for motif prediction

DESSO contains a CNN model for motif patterns learning and a novel statistical model for motif instances identification. It enables extraction of more complex motif patterns comparing with existing motif prediction methods owing to its multi-layer network architecture. We designed a first-of-its-kind binomial-based model in DESSO to identify all the significant motif instances, under the statistical hypothesis that the number of random sequence segments which contain the motif of interest in the human genome is binomially distributed [47].

The first layer of the CNN model is a convolutional layer which contains multiple convolutional filters (**Fig. 1A**). These filters were used to identify low-level features from given input (i.e., ChIP-Seq peaks). Subsequent max pooling layer and two fully connected layers were used to extract high-level abstractive features based on the output from the convolutional layer [49]. Specifically, the CNN model takes DNA sequences centered on the peaks from ChIP-Seq as input query sequences and learns motif patterns using convolutional filters (denoted as motif detectors) [39, 50]. Then, a large set of background sequences were selected from the human genome based on the genomically equivalent negative regions (GENRE) tool, considering selective sequence features, i.e. GC content and CpG frequency, and genomic features, i.e. promoter and repeat overlaps, to eliminate biases induced by these features [51]. Both the query and background sequences were then aligned as sequence matrix, where each row represents a distinct sequence. For each optimized motif detector, two motif signal matrices were derived by sliding the detector along the query sequence matrix and background sequence matrix, respectively (**Fig. 1B**). Each element of a signal matrix represents the occurrence probability of the corresponding motif detector on a sequence segment in the corresponding sequence matrix. These two motif signal matrices were then used to generate motif candidates by varying a motif instance signal cutoff in a predefined interval. For each value of the motif signal cutoff, the motif instance candidates in the query sequence matrix and background sequence matrix were obtained, and then used to calculate a *p*-value according to the binomial distribution (**Fig. 1C**).

**Figure 1.**
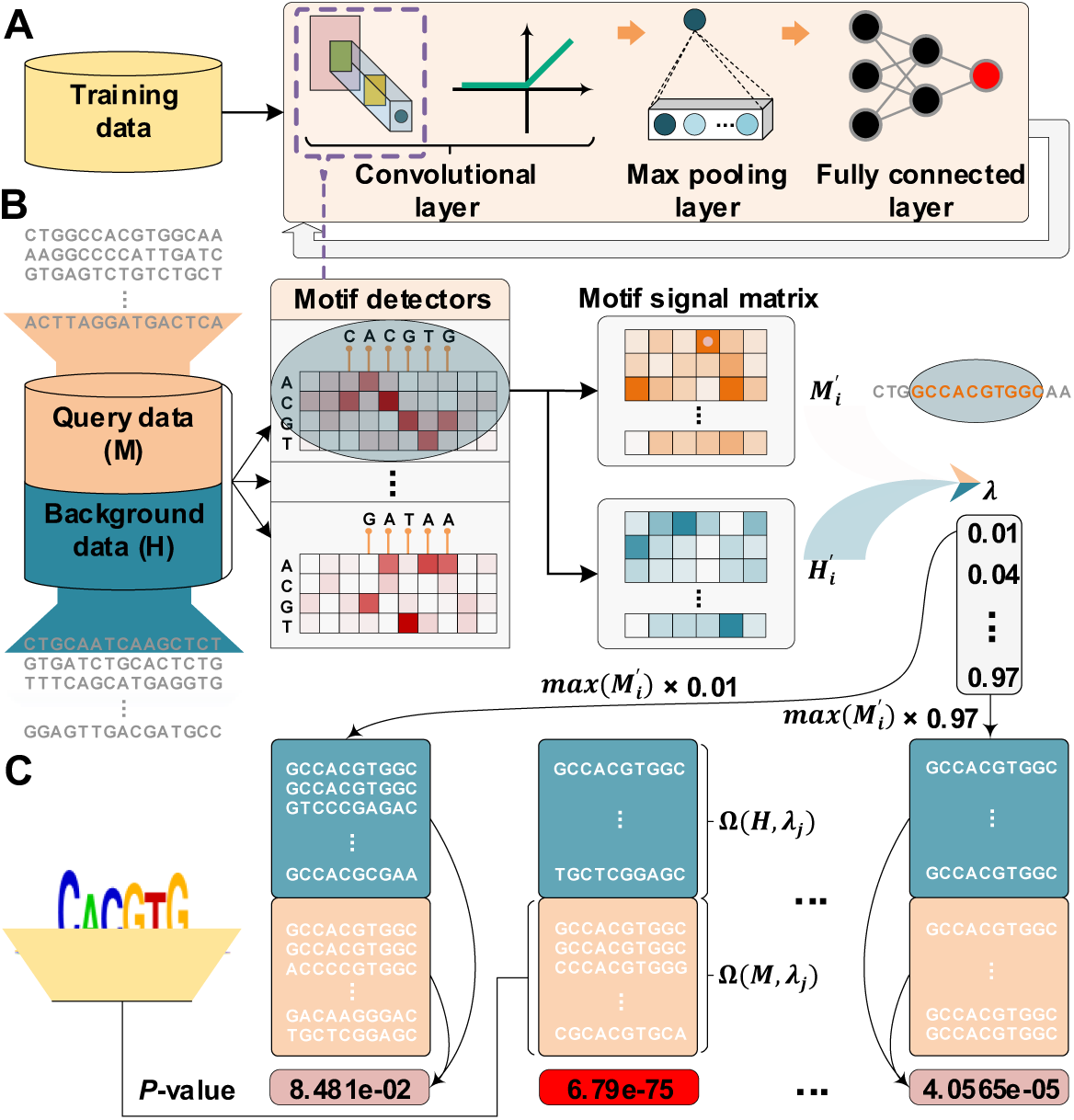
Schematic overview of DESSO framework. (**A**) The CNN model for optimizing motif detectors. (**B**) Determination of optimal motif instances recognized by each motif detector. Both the query data (*M*) from the corresponding ChIP-Seq dataset and the background data (*H*) were fed into the convolutional layer in the trained model. For each motif detector, two motif signal matrices 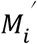 and 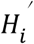 representing the probability of motif occurrence at each position in *M* and *H* were derived, respectively. (**C**) Construction of the optimized motif profile. 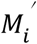 and 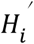> were then used to determine motif instance sets Ω(*M, λ*_*j*_) and Ω(*H, λ*_*j*_) in *M* and *H* by varying the cutoff *λ* ∈ (0, 1). For each *λ*_*j*_, a *p*-value was approximated by the binomial distribution based on the number of motif instances in Ω(*M, λ*_*j*_) and Ω(*H, λ*_*j*_). The motif instances Ω(*M, λ*_*j*_) corresponding to the minimum *p*-value were used to generate the motif logo.

The optimal motif instances for a motif detector were finally defined as the motif instance candidates in the query sequence matrix that corresponds to the minimum *p*-value (see details in the **METHODS** section). The results showed that DESSO significantly improved the motif prediction performance on 690 ENCODE TF ChIP-Seq datasets, which cover 161 TFs in 91 cell lines, in terms of the number of distinctly predicted motifs and the similarity to annotated motifs in various databases (see details in the following section).

### DESSO accurately predicts motifs from ChIP-Seq data

To test DESSO’s accuracy similarity comparisons between predicted and experimentally validated Homo sapiens motifs in the JASPAR [16] and TRANSFAC [15] databases was conducted using TOMTOM [52]; and we compared four widely used methods with DESSO, i.e., DeepBind [41], Basset [43], MEME-ChIP [35], and gkm-SVM [46] (see details in the **METHODS** section).

The numbers of distinctly validated motifs in the JASPAR and TRANSFAC covered by each of the above five methods were compared. The results showcased that DESSO achieved the best performance with 274 validated motifs covered in total, relative to 269, 269, 270, and 220 validated motifs covered by DeepBind, Basset, MEME-ChIP, and gkm-SVM, respectively (**Fig. 2A**). It is noteworthy that the number of validated motifs covered by MEME-ChIP is almost the same as the other three DL-based methods (DESSO, DeepBind, and Basset). This performance may be attributed to the fact that MEME-ChIP considered not only motifs recognized by single TF and TF complexes but also very short motifs (up to 8 bps) that preferred by most eukaryotic monomeric TFs. Five validated motifs were exclusively covered by DESSO, which are recognized by NR3C2, FOS::JUN (var. 2), RFX3, HIC2, and DUXA, respectively (**Fig. 2B**). Specifically, NR3C2 plays an essential role in mediating ion and waster transport [53], and DUXA is closely associated with Facioscapulohumeral Muscular Dystrophy 1. FOS::JUN (var. 2) is one of the JUN-FOS heterodimers [54], which was first discovered by the selective microfluidics-based ligand enrichment followed by sequencing [55].

**Figure 2.**
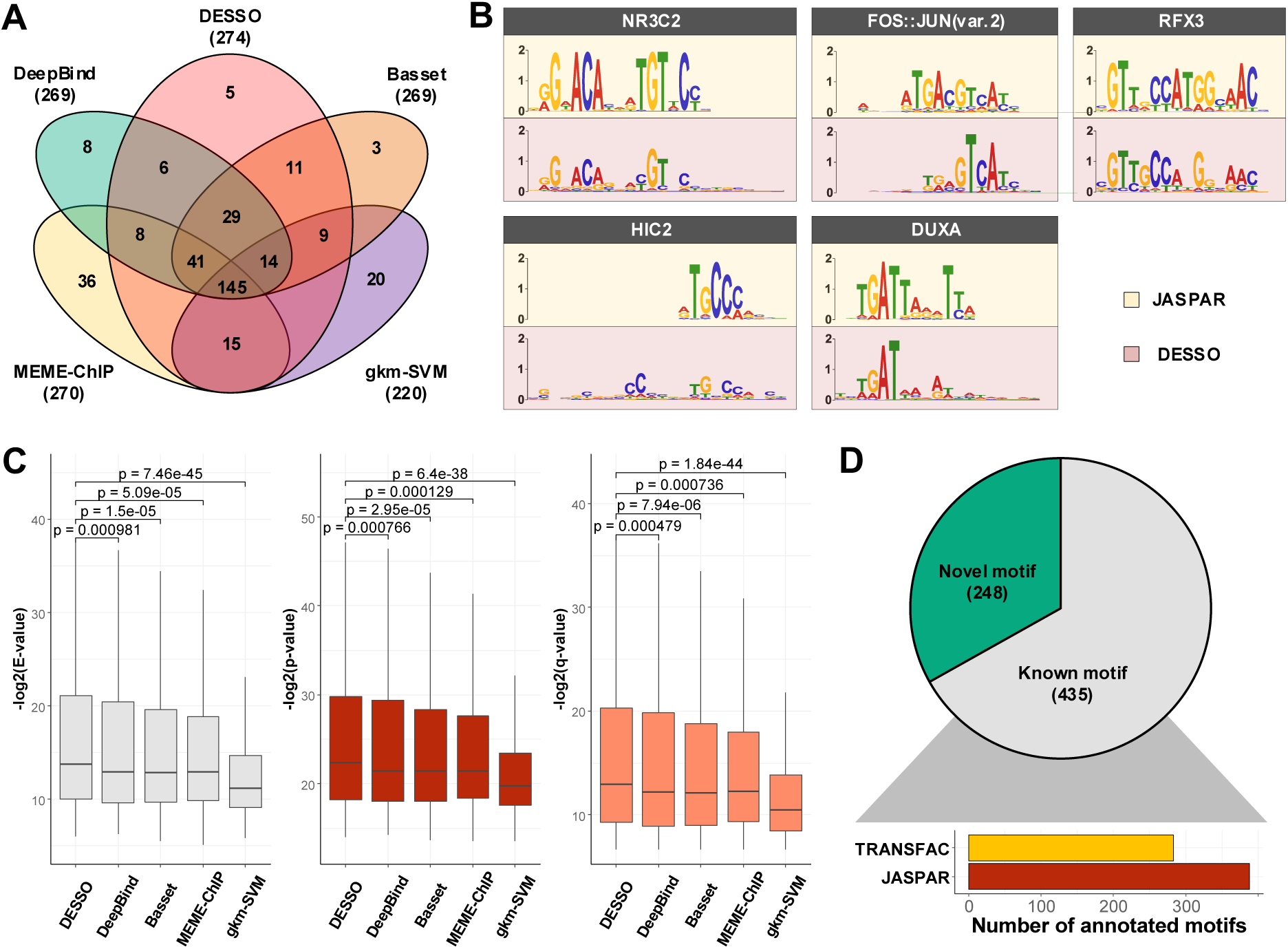
A performance comparison of sequence motif identification accuracy. (**A**) Venn diagram of the validated motifs in the JASPAR and TRANSFAC identified by the five tools. (**B**) The five validated motifs that were uniquely identified by DESSO. (**C**) The - *log*2 (E-value), - *log*2 (*p*-value), and - *log*2 (q-value) derived from TOMTOM for all methods. The Wilcoxon test *p*-values of the above three scores between DESSO and other four methods. (**D**) A total of 435 motifs (known motifs) from DESSO can be matched to the JASPAR or TRANSFAC, and 248 motifs (novel motifs) do not have any matches (pie chart). For these known motifs, 388 of them are in the JASPAR and 283 of which are in the TRANSFAC (bar plot).

To investigate the accuracy of the sequence motifs identified by DESSO, TOMTOM was used to compare the statistical significance (i.e., *E*-value, *p*-value, and *q*-value) across JASPAR and TRANSFAC for motifs that were predicted by all methods. These statistical measurements quantify the similarity of query motifs against validated motifs in a motif database. The - *log*2(*E*- value) of DESSO was significantly larger than other methods (all achieving Wilcoxon test *p*- values < 1×10^-3^, shown in **Fig. 2C**). Similar phenomena were observed for - *log*2 (*p*-value) and - *log*2 (*q*-value) (**Fig. 2C)**. DeepBind, Basset, and MEME-ChIP achieved comparable performance across the three measurements and significant performance over gkm-SVM. Thus, DL frameworks were able to learn motif patterns from DNA sequences more accurately than the combination of expectation maximization algorithm and regular expression, and support vector machine. Most importantly, these results highlight the advantage DESSO’s binomial model over strategies used by DeepBind and Basset and demonstrate that DESSO reduces both false positive and false negative rates.

To explore the sequence motifs identified by DESSO, clustering according to their similarity score from TOMTOM give rise to 683 clusters **(Fig. 2D)**. The most significant motif in terms of the binomial *p*-value in each cluster was defined as the representative sequence motif. Among the 683 representative sequence motifs, 435 of them were known motifs, found in the JASPAR or TRANSFAC, while 248 of them were novel motifs, not previously validated (**Fig. 2D** and see details in Methods) and retained for additional analysis owing to their statistical significance.

### Analysis of 683 representative sequence motifs identified by DESSO

The structural classes of human TFs or TF complexes that recognize the 435 known motifs identified by DESSO were analyzed with TFClass [56]. Twenty-six structure classes, including five dimerization structures and 21 monomer structures were represented in the identified motifs (**Fig. 3A**). The C2H2 zinc finger (C2H2) was the most prevalent structural class, representing 89 of the 435 known motifs, consistent with the fact that C2H2 represents one of the most common TF domains in humans [57]. A total of 88 representative motifs that were frequently observed in the 690 ChIP-Seq datasets, were derived from these 435 known motifs [58] based on the motif comparison tool, TOMTOM, and the hierarchical clustering (**Fig. 3B,** see details in METHODS).

**Figure 3.**
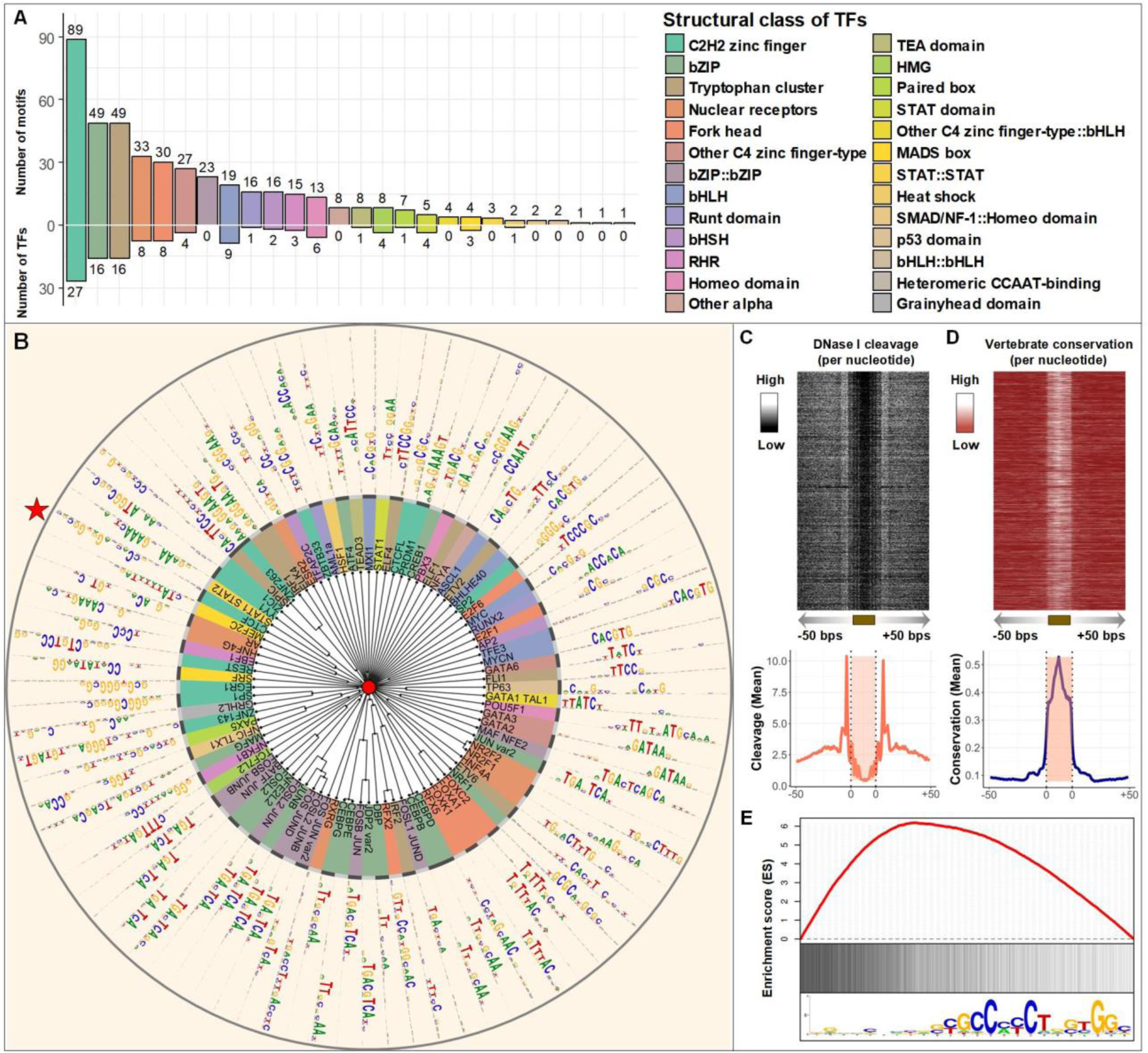
Detailed view of identified sequence motifs. (**A**) The number of know motifs identified by DESSO and the number of TFs in 690 ChIP-Seq datasets represented by the 26 structural classes in the TFClass system. (**B**) The phylogram tree of 88 enriched sequence motifs according to their similarity derived from TOMTOM. The inner circle indicates the TFs and their structural classes (background color is the same as **A**) of the corresponding validated motifs either in the JASPAR or TRANSFAC. The outer circle represents the motif logo of each enriched sequence motif, including CTCF (red star) and MYCN (green star). (**C**) The heat map of per-nucleotide DNase I cleavage and the corresponding mean value; (**D**) The heat map of per-nucleotide vertebrate conservation and the corresponding mean value of CTCF’s TFBSs as well as ±50 bps flanking regions within the A549 cell line. Each row in the heat map represents a motif instance. (**E**) Red curve indicates the enrichment score of CTCF on its corresponding ChIP-Seq peaks. Vertical black lines indicate the presence of ChIP-Seq peaks that contain at least one TFBS of CTCF. The motif logo of CTCF identified by DESSO is shown at the below.

Various functional analyses have been carried out for the 88 motifs, aiming to illustrate the overall quality of the motifs in ChIP-Seq data that DESSO identified (see details in DESSO website). Among the 88 motifs, the CTCF motif, that plays an important role in modulating chromatin structure [61], was the most enriched as it is the ChIP-ed TF for approximately 15% ChIP-Seq datasets in our study [59, 60]. Thus, we selected CTCF as the example for the following functional analyses. The DNase I Digital Genomic Footprinting [62] and evolutionary conservation (phastCons scores [63]) of CTCF’s TFBSs within A549 cell line were collected. CTCF’s TFBSs were more susceptible to DNase I enzyme (**Fig. 3C**) revealing that the binding preference of CTCF to accessible chromatin and showed significant evolutionary conservation compared to the flanking regions (**Fig. 3D**), illustrating a strong phylogenetic conservation of the identified CTCF motif. To investigate the occurrence of CTCF motif in the corresponding ChIP-Seq peaks which are ranked by peak signal, its enrichment score was calculated using GSEA (Gene Set Enrichment Analysis software) [64] (**Fig. 3E**). The enrichment score curve clearly showed the dramatic left-skewed trend, indicating that the DESSO identified CTCF motif was more enriched in top-ranked peaks. This is consistent with the fact that peaks with higher peak signal also have a higher probability to be bound by the ChIP-ed TF. In addition, another well-known TF, MAX, also demonstrated strong functionality conservation and left-skewed enrichment in K562 cell line (Supplementary **Fig. S1A-C**).

An extended investigation of the 248 novel motifs showed that they are very likely to be *cis*- regulatory elements (**Fig. S2**) but have not been experimentally validated and demonstrated having similar functionalities as known motifs. Seventy-eight distinctly enriched motifs were collected from the 248 motifs based on similar clustering analyses as above (See Supplementary **Fig. S2A** and details in Methods) [65]. The functionality and enrichment analysis of these motifs also demonstrated strong DNase footprint patterns and evolutionary conservation, revealing the potential role of this motif in transcriptional regulation (Supplementary **Fig. S2B-D**). These results strongly supported the functionality conservation of both known motifs and novel motifs identified by DESSO and proved the distinguished ability of DESSO in identifying regulatory code in the human genome.

### DESSO infers indirect binding mechanisms from the 100 TFs expressed in the K562 cell line

Increasing evidence indicates that direct TF-DNA interaction is not the only way for TFs to interact with DNA [66]. Thus, sequence motifs identified by DESSO in each of the 690 datasets may also contain motifs that were bound by other TFs that may have been associated with the motif via partner, rather than a direct interaction via the ChIP-ed TFs. To investigate this phenomenon for human TFs, the 100 TFs in the K562 cell line were examined by analyzing their corresponding sequence motifs identified by DESSO (Supplementary Table S1). Among the 100 TFs, 75 of them are DNA-binding proteins involved in RNA polymerase II transcription [67], which were referred to as the sequence-specific TFs. The remaining 25 TFs are non-sequence-specific and are thought to interact with DNA by other ways including protein-protein interactions with other DNA-binding proteins. Sixty-seven of the 75 sequence-specific TFs have known/annotated canonical motifs, which represent sequence patterns that are specifically recognized by their DNA-binding domains [68]. For each of their corresponding ChIP-Seq datasets, the peaks containing the canonical motif of the ChIP-ed TF were defined as direct-binding peaks (***D***) and the others were defined as indirect-binding peaks (***I***). Fifty-three of the 75 TFs have their canonical motifs been discovered, indicating DESSO was able to identify 80% of the canonical motifs. Approximately 48% of the ChIP-Seq peaks for these 53 TFs belong to ***I*** on average, and this proportion (72% average) was observed across all the 100 TFs (blue bars in the outer ring of **Fig. 4** [65]).

**Figure 4.**
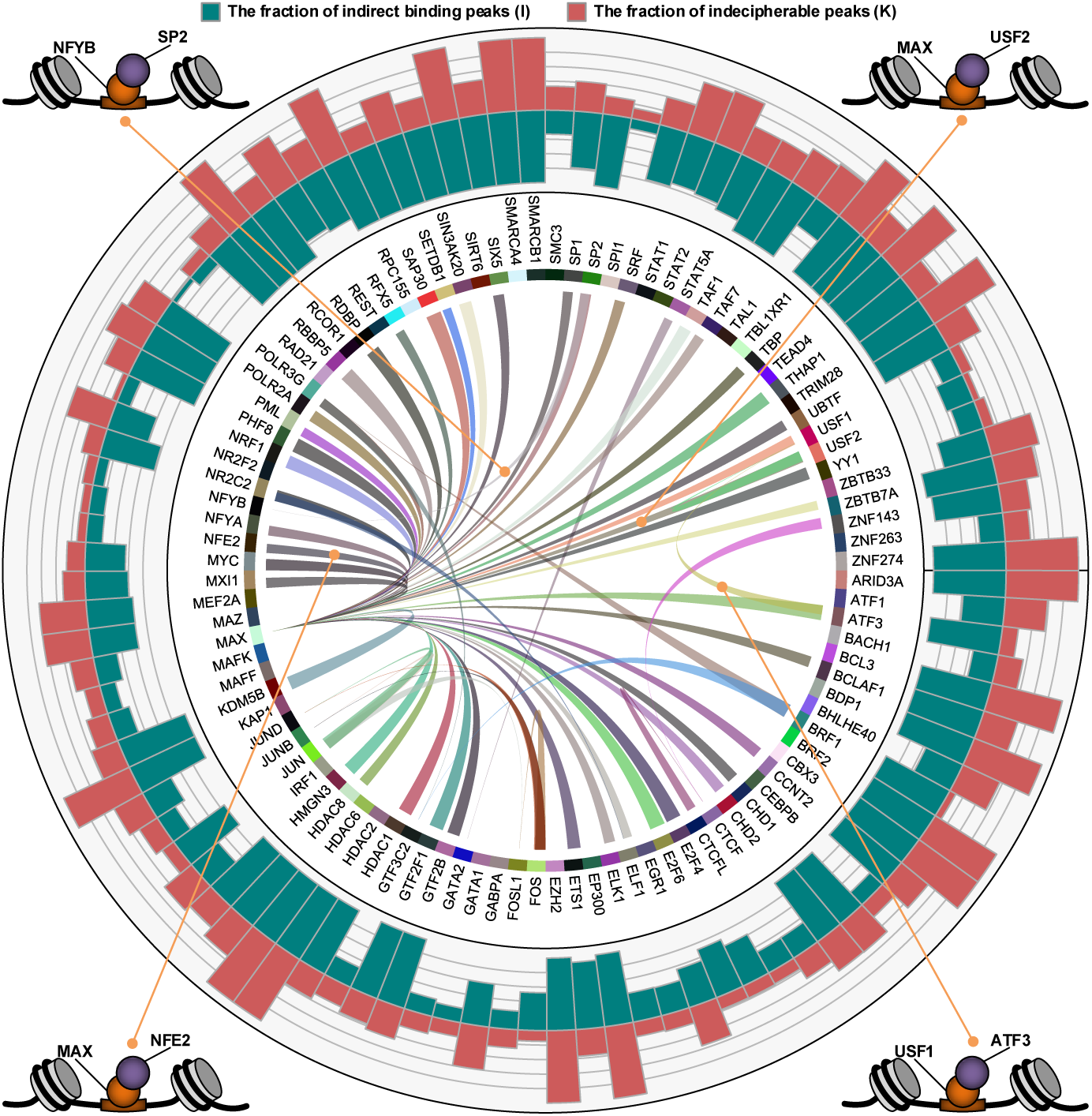
Indirect binding of the 100 TFs analyzed in the K562 cell line. The names of the 100 TFs are indicated around the inner circle, and a ribbon connects two TFs which have predicted tethering binding association. The thickness of the ribbon is proportional to the ratio of peaks in the wide-sided TF’s *I* (indirect) that are consistent with the narrow-sided TF’s *D* (direct). The blue bar and red bar in the outer circle indicate the ratio of *I* and the ratio of ***K*** (indecipherable - cannot be resolved as tethering/binding) in each TF’s ChIP-Seq dataset, respectively. Four examples of tethering binding association are showcased around the outer circle, each of which indicates that one TF (lavender ball) interacts with DNA by binding to another DNA-binding TF (orange ball).

A likely reason for this observation is that rather than bind to DNA sequence directly, some sequence-specific TFs can also tether to DNA by interacting with other DNA-binding proteins [69]. Such indirect binding is abundant in human TFs, e.g., the estrogen receptor a is enabled to regulate gene expression by interacting with Runx1 in breast cancer cells [70] and interact with c-Fos/c-Jun heterodimers at TFBSs of AP-1 in ER/AP-1-dependent transcription [71]. To further investigate DESSO’s ability to predict tethering and pairwise binding among these 100 TFs, we calculate the proportion of ***I*** peaks in one TF’s that are consistent with the ***D*** peaks of another TF (see Supplementary Table S2 and details in the **METHODS** section). A total of 61 tethering binding associations were discovered (the links in the inner ring of **Fig. 4**). These included two known tethering binding mechanisms (i.e., ATF3-USF1 [68] and NFE2-MAX [62]) and some potential interactions which have been observed in recent studies, such as USF2-MAX and SP2-NFYB [62] (in the four corners of **Fig. 4**).

Notably, our results reported that 45 TFs have tethering interactions with MAX, of which, 30 belonged to TFs that had sequence-specific motifs and the remaining 15 were non-sequence-specific TFs (Supplementary Table S1). Out of these 45 tethering interactions, 7 (15.6%) of them were validated interactors with MAX by the PPIs in the BIOGRID database and were documented MAX-associated binders [72]. As a basic helix-loop-helix zipper (bHLHZ) TF, the biological function of MAX [73] can only be activated by forming dimers/complexes with other proteins. Importantly, MAX was always the DNA binder reinforcing the idea that it serves as the tethering sites for many other TFs. The most well-known MAX-associated complex is the MYC/MAX/MAD network, including MYC-MAX and MAD-MAX heterodimers which are widely recognized to play an important role in cell proliferation, differentiation, and neoplastic disease [74, 75]. Our observation revealed that not only specifically dimerizing with proteins in MYC family [73], MAX also extensively interact with other sequence-specific and non-sequence-specific TFs from diverse protein families.

For each ChIP-Seq dataset of the 100 TFs, the peaks in ***I*** that do not involve in any tethering binding interactions were classified as indecipherable peaks (***K***), indicating the peaks that cannot be deciphered based on direct DNA binding and tethering binding mechanisms. These peaks composed about 49% of all ChIP-Seq peaks in these 100 datasets (red bars in the outer ring of **Fig. 4**). Furthermore, even no any statistically significant sequence motifs were identified by DESSO in a total of 51 of the 690 ChIP-Seq datasets. Taken together, these analyses implied that sequence motifs still have considerable limitations in elucidating TF-DNA recognition in human, so advanced mechanisms which may occur even beyond sequence-level TF-DNA interactions should be considered. An emerging feature for elucidating the advanced mechanisms is DNA shape, which will be detailed analyzed in the following two sections.

### DESSO recognized DNA shape features as contributors to TF-DNA binding specificity

To investigate the importance of DNA structure in human TF-DNA recognition, DESSO was used to infer the power of DNA shape in predicting TF-DNA binding specificity across the 690 ChIP-Seq datasets. For each dataset, the 101-bp sequences centered at their peak summits were defined as positive sequences [41]. Additionally, the corresponding negative sequences were selected from the human genome provided they do not have any overlaps with peaks in that dataset and they have the same GC-content as the positive sequences. The HelT, MGW, ProT, and Roll of each positive and negative sequence were then generated by DNAshapeR [76] and used to train DESSO.

DESSO was then applied to HelT, MGW, ProT, Roll, and the combination of these four shape features (referred to as *DNA shape combination*) to classify the positive and negative sequences in each dataset (**Fig. 5A** and **Supplementary Fig. S3**). For the five kinds of inputs, their performance was evaluated using the area under the receiver operating characteristic curve (AUC). Specifically, HelT, MGW, ProT, and Roll achieved an average AUC of 0.72, 0.66, 0.69, and 0.75, respectively (**Fig. 5B**). It is clear that HelT and Roll substantially surpassed the classification performance of MGW and ProT, which may stem from the fact that HelT and Roll were calculated by the two central bp steps within a sliding-pentamer window (secondary structure information), while MGW and ProT were calculated by only the central bp [26]. Hence DESSO predicts that DNA shape factors HelT and Roll have significant predictive power in identifying TFBS.

**Figure 5.**
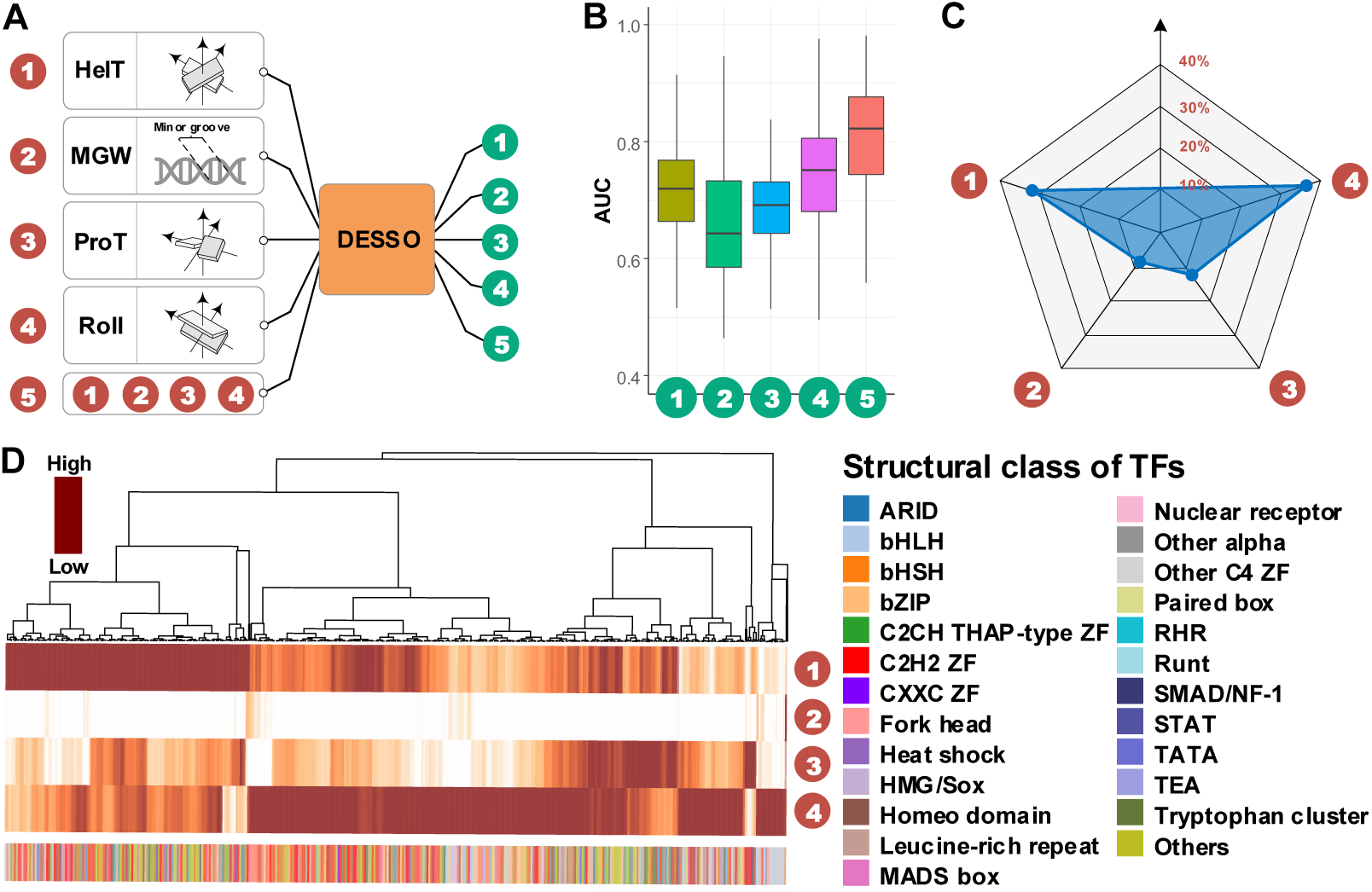
The performance of DNA shape in predicting TF-DNA binding specificity. (**A**) DESSO was applied to five different inputs, i.e., HelT, MGW, ProT, Roll, and DNA shape combination. (**B**) The average AUC of the five inputs based on the 690 ChIP-Seq datasets. (**C**) The contribution of HelT (32%), MGW (9%), ProT (22%), and Roll (37%) in DNA shape combination in predicting TF-DNA binding specificity. (**D**) The heat map is a more detailed analysis of diagram **C**, indicating the contribution of each DNA shape feature on the 690 datasets, where each column represents a dataset. Those columns were organized by hierarchical clustering based on Pearson correlation and complete linkage. The structure class of ChIP-ed TF in each dataset was showcased at the bottom.

Compared to individual DNA shape features, the performance of DNA shape combination was significantly improved (AUC of 0.81 average) (**Fig. 5B**), indicating the complementary role of HelT, MGW, ProT, and Roll in predicting TF-DNA binding specificity. To evaluate to what extent each DNA shape feature quantitatively contribute to such remarkable performance, the fraction of the average motif signal from the max pooling layer for each kind of shape feature was calculated for the 690 datasets. The results reported that HelT, MGW, ProT, and Roll contributed 32%, 9%, 22%, and 37%, respectively (**Fig. 5C**). Thus, the prediction from the DESSO analysis indicated that the shape factors HelT, ProT and Roll frequently contribute simultaneously to TF binding. To assess the common occurrences of DNA shape factors across individuals from the 690 datasets (with DNA binding domain information of the ChIP-ed TF available), we analyzed the DESSO results and clustered them (**Fig. 5D**). These results indicated that HelT and Roll were the most important contributors and that ProT appeared predicative within only for a small clave of samples. Thus, DNA shape factors tended to have dominant roles in different datasets (Fig. 5D). Surprisingly, the ChIP-ed TFs had no apparent predictive power on the dominant shape factor suggesting that there may be more rules or additional shape factor information to be uncovered.

Overall, these results demonstrated the remarkable predictive performance of DNA shape features in TF-DNA binding specificity prediction, implying that the underlying conserved DNA shape patterns (or *shape motif*) are also encoded in the human genome and may involve in TF-shape readouts recognition. More details of the shape motifs analyses can be found in the following section.

### DESSO predicts novel DNA shape motifs

We applied DESSO to determine if the human genome contained regions of evolutionarily conserved shape motifs. Specifically, we sought to have DESSO discover shape motifs based on the same strategy that was used to discover sequence motifs (similar to Fig. 1). This approach added HelT, MGW, ProT, Roll or their combinations to discover four kinds of shape motifs within the 690 datasets from ENCODE (named HelT motif, MGW motif, ProT motif, and Roll motif). DESSO identified 1,257 HelT motifs, 84 MGW motifs, 885 ProT motifs, and 478 Roll motifs, with 598 out of the 690 datasets having at least one shape motif. A shape motif can be represented by a vector of shape features describing the mean of the corresponding motif instances. Using the same strategy as in figure 5D, and counting the shape motifs belonging to each dataset, we found that the distribution of shape motifs across the datasets were enriched for HelT motifs and ProT motifs (**Fig. 6A**). It was surprising that given the large fraction of Roll shape factors and low abundance of ProT shape factors (Fig. 5C and 5D), that a larger number of shape motifs was identified for ProT (Fig 6A). Overall, these results indicate that DESSO was able to identify shape motifs across a large range of datasets indicating that shape motifs are abundant in the human genome.

**Figure 6.**
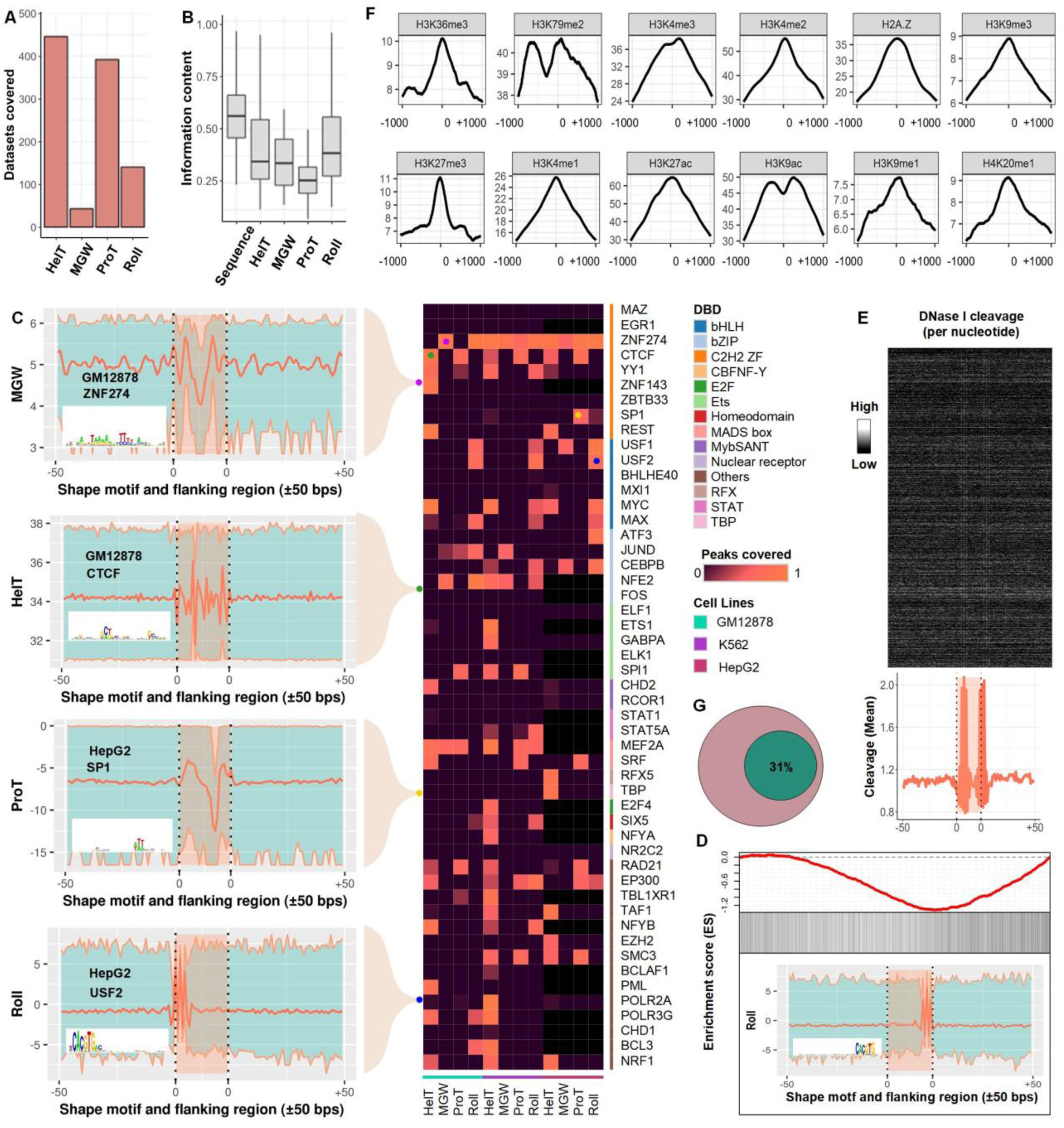
A comprehensive analysis of the identified shape motifs. (**A**) The number of datasets in the 690 ENCODE ChIP-Seq datasets that were covered by shape motifs of HelT (446), MGW (43), ProT (392), and Roll (141). (**B**) IC of underlying sequences of the identified sequence motifs and shape motifs. (**C**) Each entry in the heatmap indicates the ratio of peaks covered by the four kinds of shape motifs which were identified in the 51 TFs within GM12878, K562, and HepG2 cell lines, while the black entries represent missing values. Four representative shape motif logos were listed at the left side, where each of them represents the shape motif profile and ±50 bps flanking regions using a bold orange curve. The two boundary curves of the blue region represent upper and lower bounds of shape features in the corresponding motif instances. The logo of shape-sequence-motif is showcased at the lower-left corner of each shape motif logo. (**D**) The enrichment score of MAX’s Roll motif in its corresponding ChIP-Seq peaks in the K562 cell line. Black ticks indicate the occurrence of ChIP-Seq peaks that contain at least one instances of MAX’s Roll motif. The logo of MAX’s Roll motif is shown at the below. (**E**) The heat map of per-nucleotide DNase I cleavage of TFBSs of MAX’s Roll as well as ±50 bps flanking regions, where each row represents a motif instance. The orange curve represents mean DNase I cleavage. (**F**) Twelve histone marks of ±1000 bps around the summits of Max’s Roll motif. (**G**) The average ratio (31%) of ChIP-Seq peaks that cannot be explained of the 100 TFs within K562 cell line.

Given that the shape features were derived from conserved DNA sequences, we predicted that the newly identified shape motifs should have a high probability of coinciding with shape features within the sequence motifs in their respective datasets. To examine this hypothesis, the underlying DNA sequences of each shape motif were aligned as a sequence motif profile which we defined as a shape-sequence-motif. The information content (IC) of each shape motif class was then computed across the shape-sequence-motifs (**Fig. 6B**). Compared with the sequence motifs identified by DESSO (Fig. 1), shape-sequence-motifs have significantly lower IC (**Fig. 6B**). We also measured the similarity between each shape-sequence-motif and validated motifs in JASPAR and TRANSFAC using TOMTOM. Only 66% of shape-sequence-motifs can be matched to the JASPAR or TRANSFAC. Taken together, these two results indicate that shape motifs are less conserved at the sequence level and are largely independent from sequence motifs.

To investigate the enrichment of shape motifs and whether they are cell-line-specific or DNA-binding-domain-specific, the fraction of peaks that were covered by each kind of shape motifs of the 51 TFs within GM12878, K562, and HepG2 cell line were analyzed. The majority of peaks in ZNF274’s datasets can be accounted for by its shape motifs (**Fig. 6C**), even though no peaks were explained by direct TF-DNA binding and tethering (**Fig. 4**). CTCF coherently recognizes HelT and ProT motifs among the three aforementioned cell lines, while SP1 is dominated explicitly by ProT motifs within the HepG2 cell line. Also, the Roll motif is prevalent in TFs which have basic helix-loop-helix (bHLH) structure, implying that Roll is recognized explicitly by such a DNA binding domain (**Fig. 6C**). To explore the occurrence of shape motifs in their corresponding ChIP-Seq peaks, enrichment analysis of Max’s Roll motif in the K562 cell line was performed. Opposed to its sequence motif, Max’s Roll motif is more enriched in the low-ranked peaks which is consistent with the recent observation that TFs bind to peaks with low peak signal by recognizing their preferred shape profiles [24] (**Fig. 6D**). We then examined whether Max’s Roll motif is functionally conserved by analyzing DNase I Digital Genomic Footprinting and 12 histone marks surrounding its TFBSs [77] and found that Max’s Roll motif also prefers histone-depletion regions like its sequence motifs (**Fig. 6E-F**).

Considering ChIP-Seq peaks covered by shape motifs, the average ratio of ***K*** of the 100 TFs within the K562 cell line (Fig. 4) was decreased to 31% (**Fig. 6G**). This result suggested that human TFs are capable of recognizing shape motifs in the genome, which contributes to explaining the ChIP-Seq peaks that cannot be interpreted by direct TF-DNA interaction and tethering binding.

### DESSO web server

To broadly facilitate motif-related analysis in this field, we also provide an integrated web server for DESSO, which is freely available at http://bmbl.sdstate.edu/DESSO. In addition to showing identified sequence and shape motifs from the 690 ENCODE ChIP-Seq datasets, functionally conserved analyses of each motif were also provided on this web server. Furthermore, DESSO enables motif scan and other comprehensive analyses based on user-provided DNA sequences. The source code of DESSO and a detailed tutorial can be found at https://github.com/viyjy/DESSO.

## CONCLUSION AND DISCUSSION

We developed a DL-based motif finding framework, DESSO, combed with a new statistical method for motif profile construction, followed by its application on the 690 human ChIP-Seq datasets within ENCODE. This work lead to a first ever, comprehensive analyses of identified sequence and shape motifs. DESSO improved the state-of-the-art performance of *cis*-regulatory motif prediction and TFBSs identification and showcased the potential of a DL framework for identification and rationalization of results. Our results demonstrate that DESSO was able to identify a number of previously unidentified motifs and shape factors that contribute to TF-DNA binding mechanisms and can infer the indirect regulation mechanisms through tethering binding activities and co-factor motifs predictions. These predictions now await experimental validation. Overall, the implementation and application of the framework provide a solid foundation for the construction of gene regulatory network and the elucidation of TF-DNA binding mechanism in the human genome.

An important area in this filed is how motif flanking regions influence TF-DNA binding [78-80]. Although most existing studies focused on proximal flanking regions around core motifs, it is still not clear whether more remote flanking regions will facilitate TF-DNA binding. Further investigations were carried out in this study using the 181 ChIP-Seq datasets with predicted AUC values of less than 0.9 in DeepBind. The results showcased that the test AUC increases with increasing peak length (**Figure 7A**). Specifically, increasing peak length to 1,001 bps results in an AUC of 0.98 on average. This implies that the flanking regions of ChIP-Seq peaks contain useful information in TF binding, and hence, using flanking regions as negative sequences is inappropriate [47, 81, 82].

**Figure 7.**
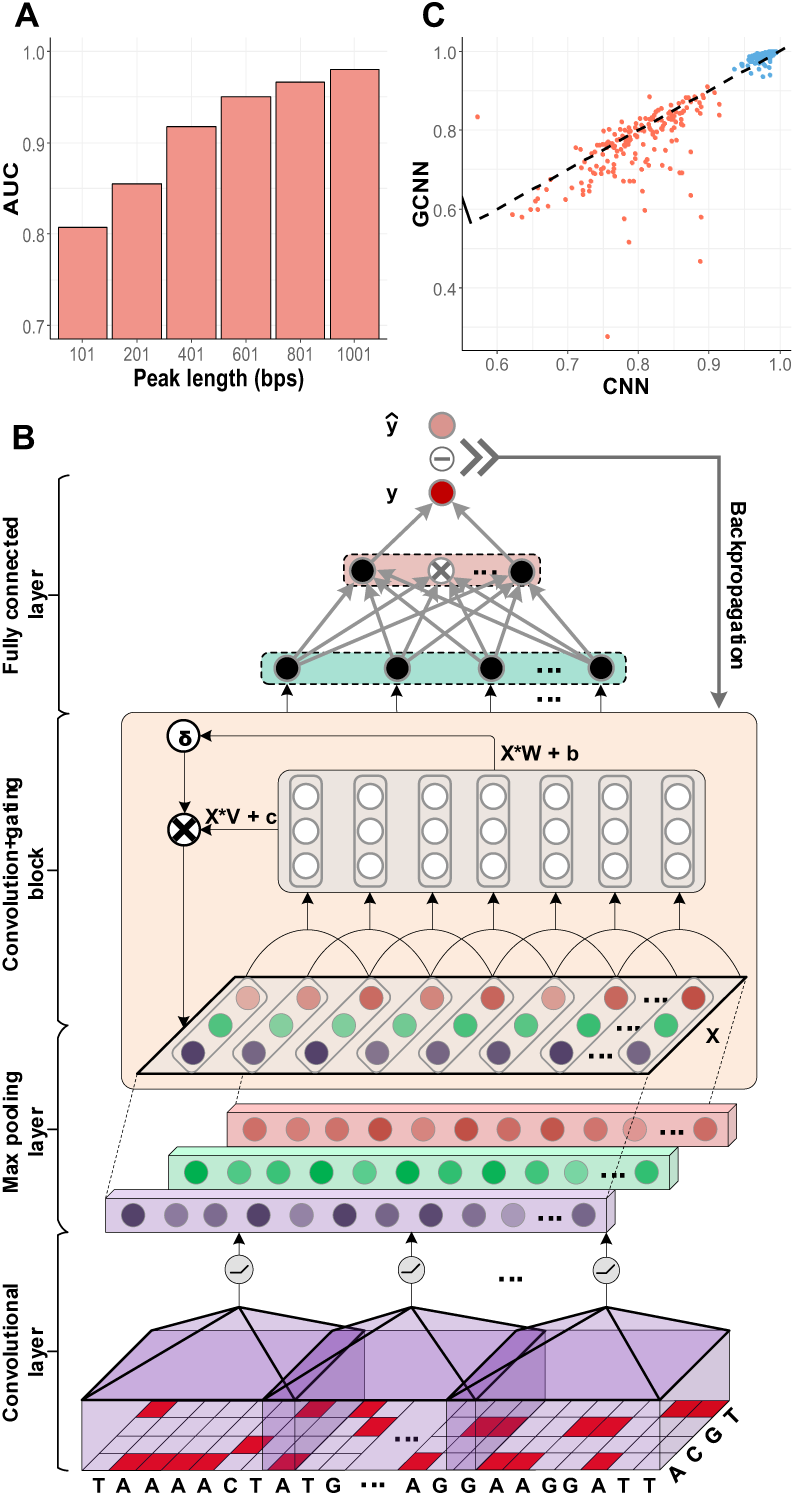
The application of GCNN in TF-DNA binding specificity prediction. (**A**) As peak length increases, the predictive performance of CNN on the 181 ChIP-Seq datasets was improved monotonically. (**B**) The workflow of GCNN with DNA sequence as input. (**C**) The AUC of GCNN and CNN on the 181 datasets based on the peaks with 101 bps length (orange points) and 1,001 bps length (blue points).

Although CNN enables more complex motif pattern extraction owing to its multi-layer architecture, it is limited in the ability to capture the long-range dependencies among motifs. Inspired by the recurrent neural network (RNN), which can capture the unbounded context in natural language, the models combining CNN and RNN (CNN-RNN) have significantly improved prediction of TF-DNA binding specificity [44, 83]. The downside of RNN is its inability to parallelize over sequential inputs, resulting in substantial processing steps as the length of the input increases. Alternatively, the gated convolutional neural network (GCNN) has been proposed recently and performed competitively on benchmarks [84]. It allows parallelization by stacking convolutions but still has the capability in capturing long-range dependencies of inputs. We proposed a GCNN-based model for long DNA sequences (1,001bp) in TF-DNA binding specificity prediction (**Figure 7B**). This model contains a convolutional layer, a recurrent convolution-gating block (CGB), and two fully connected layers. Concretely, the convolutional layer aims to detect motifs, the CGB captures long-term dependencies among identified motifs, and the fully connected layers account for TF-DNA binding specificity prediction. Among the 181 datasets mentioned above, our GCNN model achieved higher AUC than CNN on 169 of them (**Figure 7C**). This remarkable performance benefits from its ability in capturing long-term dependencies among identified motifs (not necessary the *cis*-regulatory motifs). With a competitive performance compared with CNN-RNN, GCNN is more efficient in the application on genome-scale, because it allows parallelization by stacking convolutions thus enhanced the ability to process high throughput data set.

Indeed, further investigations are needed to elucidate other obscure intrinsic features in gene regulation and TF binding. Specifically, in this study, 31% of the peaks in ChIP-Seq data remain unexplained by sequence and shape motifs. Gene expression is controlled by multiple underlying transcriptional regulatory signals, which is a complicated process requiring the coordination of histone modifications, TF binding, and other chromatin remodeling activities. The epigenetic information as measured by histone modifications, DNase-seq, ATAC-seq, and even gene expression data have been used to predict the binding of DNA regulatory elements [85, 86]. Recently, the matched expression data and access data across diverse cellular contexts were integrated into a model to predict the missing parts, including TF binding location, chromatin accessibility, and gene expression [87]. A potential improvement lies in the evaluation of TF-DNA binding relationship, which usually conducted by calculating the significance of candidate sites based on motif occurrence frequency models including PWM. However, the frequent occurrence of motif patterns does not naturally mean high binding strength and high regulation affection.

The DL-based models provide a promising opportunity to describe the relationships between motifs and expression accurately. We believe that the accuracy of predicted motifs, deep analysis on both sequence and shape motifs and followed future studies on this research line will facilitate inference of gene regulatory relations, and accurate modeling of the complex regulatory system in the human genome. Advanced mathematics and computational tools will permit the building of integrated models of gene regulatory systems and enable deliverable strategies to prevent or treat disease.

## METHODS

### Data acquisition

The 690 ChIP-Seq datasets of uniform TFBS based on March 2012 ENCODE data freeze were downloaded from the ENCODE Analysis Database at UCSC (https://genome.ucsc.edu/ENCODE/downloads.html). These datasets contained 161 TFs and cover 91 human cell types [48]. Each dataset contained a number of peaks (ranging from 101 to 92,358), ranked in the decreasing order of their signal scores. These peaks were derived from the SPP peak caller [88] and de-noised by the Irreproducible Discovery Rate [89] based on signal reproducibility among biological replicates. The average length of the de-noised peaks is 300 bps.

A total of 53 DNase I Digital Genomic Footprinting (DNase-DGF) datasets were downloaded from the NCBI Gene Expression Omnibus (GEO) data repository (GSE26328) [62]. They provide the footprint landscape of human genome for different cell lines using the deep sequencing technique, which is based on the fact that unbound regions of regulatory factors in nucleosome-depleted chromatin are more sensitive to cleavage of DNase I.

The Vertebrate Multiz Alignment & Conservation (100 Species) by PhastCons was downloaded from UCSC (http://genome.ucsc.edu/cgi-bin/hgTrackUi?db=hg19&g=cons100way) [63]. This dataset measures evolutionary conservation of 100 vertebrate species using PhastCons based on a probabilistic model.

### DNA shape feature generation

DNA shape features (i.e., HelT, MGW, ProT, and Roll) provide three-dimensional structure information of the corresponding DNA sequences and play an essential role in TF-DNA recognition [19]. Such features were obtained by the Monte Carlo simulation and can be applied to any given nucleotide sequences by a sliding-window method [26]. A recent method designed for DNA shape analysis and its R implementation, DNAshapeR, was used to generate DNA shape features of each of the query/positive and background/negative sequences [76, 90]. All the resulting feature vectors were normalized to [0, 1].

### Overall design of DESSO

The DESSO framework is composed of (i) a CNN model for extracting motif patterns from given ChIP-Seq peaks, and (ii) a statistical model based on the binomial distribution for optimizing motif instances identification. This framework can accept both DNA sequences and DNA shape feature as input to identify sequence and shape motifs, respectively.

### CNN model construction

The CNN model contains a convolutional layer, a max pooling layer, and two fully connected layers. As this model requires binary vectors as input, each input DNA sequence was first converted to a *n* × 4 matrix *S* in one-hot format with A = [1, 0, 0, 0], T = [0, 1, 0, 0], G = [0, 0, 1, 0], and C = [0, 0, 0, 1], where *n* = 101 [41]. This was sufficient for the convolution filters to operate on sequence (S) alone. To incorporate DNA shape into DESSO, shape matrices of each DNA sequence were generated and represented by *H* (HelT), *M* (MGW), *P* (ProT), and *S* (Roll). The input of this CNN model could be (i) *S*, (ii) *H*, (iii) *M*, (iv) *P*, (v) *R*, and (vi) [*H,M,P,R*]. Each input was first fed into the convolutional layer to get the activation score of each convolutional filter:

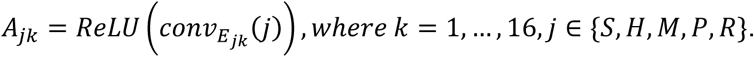

Here, the *conv*_*α*_ (*β*) represents convolution between *α* and *β, k* indicates the number of convolutional filters, and *j* indicates different input format. *E*_*Sk*_ represents convolutional filters corresponding to *S*, each of which is an *l* × 4 weight matrix with *l* = 24. *E*_*Hk*_, *E*_*Mk*_, *E*_*Pk*_, and *E*_*Rk*_ indicate convolutional filters corresponding to H, M, P, and 11, respectively, each of which is an *l* × 1 weight matrix. The *ReLU*(*x*) indicates rectified linear unit, which is a widely-used activation function in DL.

The max pooling layer was then used to downsample the activation score vectors by selecting the maximum value in each *A*_*jk*_ for *k* = 1,…,16 and *j* ∈ {*S,H,M,P,R*}. The concatenation of the output from the max pooling layer was represented by ε and finally fed into two fully connected layers. Each layer has 32 hidden neurons and used the *ReLU* activation function as above. The output layer containing only one neuron was used to predict the TF-DNA binding specificity which ranges from 0 to 1:

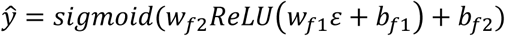

where *w*_*f*1_and *w*_*f*2_ represent the weights, while *b*_*f*1_ and *b*_*f*2_ indicate the bias units in the fully connected layers. The *sigmoid*(*x*) is a sigmoid function, where 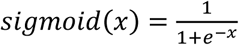.

### CNN model training

The same strategy in DeepBind [41] was used to split the peaks in each ChIP-Seq dataset into training data and test data in this study. Specifically, for a ChIP-Seq dataset, the 101-bp-long sequences centered on each peak summit was defined as positive sequences, each of which has a label of “1”. To overcome overfitting problems in the model training, for those datasets with less than 10,000 peaks, we generated complementary random peaks until having 10,000 sequences. Unlike previous studies that used dinucleotide-preserving shuffled sequences [41], regions near the transcription start sites (TSSs) [47], or flanking regions of ChIP-Seq peaks [81, 82], we picked the same number of 101-bp-long genomic sequences with same GC content from the GENCODE to generate negative sequences [91]. These negative sequences have matched GC-content to positive sequences and did not have overlap with any peaks in that dataset. Each negative sequence was labeled as “0”, which have less chance to be bound by the target TF.

For each dataset, its training data was then used to train the CNN model by minimizing the following loss function:

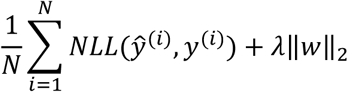

where *N* is the size of the sample set from training data (i.e., the number of sequences in the training set), *NLL*(*ŷ*^(*i*)^,*y*^(*i*)^) is negative log-likelihood between prediction *ŷ*^(*i*)^ and target *y*^(*i*)^, *λ* is a regularization parameter to leverage the trade-off between the goal of fitting and the goal of the generalizability of the trained model, and ‖. ‖_2_ indicates the *L*_2_ norm.

The loss function was optimized by mini-batch gradient descent with momentum, using a comparatively small batch size 64 to avoid the generalization drop of the trained models compared to larger batch size [92]. The backpropagation algorithm was used for gradient calculating [93], and exponential decay was applied to the learning rate with a decay rate equals 0.95. The learning rate, dropout rate, momentum, regularization parameter, and the standard deviation of the initial weights in the neural network were randomly selected from pre-prepared intervals [94]. These hyper-parameters were sampled ten times, and three-fold cross-validation was performed on the training data to select the best hyper-parameter set which corresponds to the highest average AUC. The optimal hyper-parameter set was then applied to the whole training data for the final model training. Each model mentioned above was trained for 30 epochs maximally, and an early-stopping strategy was applied to prevent overfitting. The training process was implemented based on TensorFlow which is the most widely used DL framework in public domain [95].

### Sequence motif prediction

Without loss of generality, the above CNN models, trained using DNA sequences, were used as an example to illustrate how to predict motifs based on our statistical model. Let *M* represents the 101-bp-long sequences from top-*m* peaks in each dataset, where each sequence is centered at its corresponding peak summit and *m* = min (500, *the tota1 number of peaks*). Define the motif signal matrix 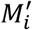 as the activation values between a motif detector *d*_*i*_ (each has length *L*=24) and *M* by feeding *M* into the convolutional layer in its corresponding trained model, and *A*_*i*_ the maximum value in 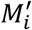. A sequence segment (*L* bps) in each sequence is defined as an *activation* segment if its activation score is larger than an activation cutoff *θ*. A motif instance set, denoted as Ω(*M, λ*), is all activation segments with *θ* = *λ* · *A*_*i*_ in *M*, where *λ* is a parameter ranging from 0 to 1. The value of *λ* could be determined by a *p*-value strategy based on the assumption that the number of activation segment containing sequences using random selection with replacement in the human genome follows a binomial distribution. To estimate the “success” probability *p* of each random selection, the human genome was divided into non-overlapping bins with length 101bp, and *n* = 500,000 bins were randomly selected as a background sequence set *H*.

Let *X* be a random variable representing the number of activation segment containing bins with *θ* = *λ* · *A*_*i*_ in *H*, *f*(*x*) = *P*(*X* = *x*) be the probability function, and *F*(*t*) = P(*X* ≥ *t*) be the cumulative distribution function. It is assumed that *f*(*x*) can be approximated by a binomial distribution *X*∼*Binomial*(*n, p*), where 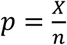 is a maximum likelihood estimate. Therefore, the *p*- value of Ω(*M*, *λ*) is given by:

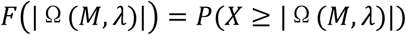

For each motif detector *d*_*i*_, the optimal motif instance Ω(*M*, *λ*)_*i*_ = *argmin*_0<*λ*<1_*F*(| Ω(*M, λ*) |) and the corresponding *p*-value can be obtained. Only Ω(*M, λ*)_*i*_ with the *p*-value less than 1 × 10^-4^, |Ω(*M, λ*)| > 5, and with at least three positions having information content larger than 1 were retained in our study, which assumes that motif should be conserved and observed more frequently in *M*. The derived motif instances were aligned as motif profiles and visualized using WebLogo 2.8.2 [96].

### Four widely used sequence motif finding tools

DeepBind [41] and Basset [43] are designed based on the same CNN model as DESSO and used the same motif detectors, query sequences, and the corresponding motif signal matrices from DESSO to learn motif patterns using motif detectors. However, they fail to optimize motif signal cutoffs for motif instance identification. For each motif detector, sequence fragments which have maximum activation score in each query sequence were aligned to obtain motifs in DeepBind, while Basset aligned sequence fragments were having activation score larger than half of *A*_*i*_ as motifs. As a highly cited web service in this field, MEME-ChIP identifies motifs from ChIP-Seq peaks by integrating two complementary motif discovery algorithms, i.e., MEME [10] and DREME [34]. Gkm-SVM was selected in the comparison as it significantly outperforms traditional kmer-SVM methods by using gapped *k*-mers for accurately and efficiently identifying longer motifs which are hard to model as *k*-mers. Specifically, The C++ implementation, gkm-SVM-2.0, was used in this study [46]. DESSO, DeepBind, and Basset were evaluated on all peaks in each ChIP-Seq dataset. Limited by the computational complexity [35], only the top 500 and top 10,000 peaks were used in MEME-ChIP and gkm-SVM, respectively, and the corresponding negative sequences used in DESSO were applied to generate sequence motifs. Besides the maximum motif length and the maximum number of output motifs were set to 24 and 16, respectively, default parameters in MEME-ChIP and gkm-SVM were used.

### Sequence motif analyses

For each of the sequence motifs identified by DESSO, it was retained for the following analysis, only if it (*i*) is more enriched in positive sequences, (*ii*) has at least five motif instances, and (*iii*) has a *p*-value less than 1*e* - 4.

All the sequence motifs, identified by DESSO, DeepBind, Basset, MEME-ChIP, and gkm-SVM, were compared with the documented Homo sapiens motifs in the JASPAR (537 motifs) [16], TRANSFAC (208 motifs) [15], and HOCOMOCO (641 motifs) [97], using TOMTOM [52] with the significance threshold FDR < 0.05. Each query motif may have multiple matched target motifs in the above databases, along with multiple comparisons provided by TOMTOM, which were listed in descending order by the similarity between the query motif and target motif. In this case, only the first comparison was used in our study. Meanwhile, the TOMTOM program generated three statistical significances for each comparison, i.e., *E*-value, *p*-value, and *q*-value.

Specifically, a total of 274 experimentally validated motifs in the JASPAR and TRANSFAC were covered by DESSO (with a TOMTOM *E*-value < 0.01), while 269, 269, 270, and 220 motifs were covered by DeepBind, Basset, MEME-ChIP, and gkm-SVM, respectively. To rule out the influence of motif detectors that are more enriched in background sequences, here, only the motif detectors complying with the above three requirements were used for motif generation in DeepBind and Basset. Furthermore, we compared the - *log*2(*E*-value), - *log* 2(*q*-value), and - *log* 2(*p*-value) of DESSO with the other four tools based on the motifs identified by all the five tools (TOMTOM *q*-value < 0.01).

Totally, 2,786 sequence motifs were predicted by DESSO (with at least three nucleotides having information content larger than 1), among which 2,263 motifs can be matched with the documented motifs in JASPAR or TRANSFAC and 523 motifs do not have any matches in these two databases. For the 2,263 motifs, 435 clusters were identified based on TOMTOM and the hierarchical clustering. For each cluster, the most significant motif was selected according to its *p*-value, giving rise to 435 motifs (http://bmbl.sdstate.edu/DESSO/knownMotif.php), where 88 of them corresponds to clusters that have greater than and equal to 3 motifs. The same procedure generated 248 clusters from the 523 motifs, in which 78 clusters have more than one motif and 178 clusters have only one motif. For each cluster, the most significant motif was selected based on its *p*-value, giving rise to 248 motifs (http://bmbl.sdstate.edu/DESSO/novelMotif.php).

### DNase I cleavage, vertebrate conservation, and enrichment analysis of identified motifs

For each identified motif, all its motif instances in the corresponding ChIP-Seq peaks were ranked by their activation score. After that, the DNase I cleavage of each motif instance along with ±50 bps flanking regions was collected from DNase-DGF [62] if the corresponding cell line is available. The evolutionary conservation of each motif instance was measured by PhastCons [63]. GSEA was used to generate the enrichment plot of each motif on the corresponding ranked peaks [64].

### Motif scan

Similar to the strategy mentioned in the ***Sequence motif prediction*** section, we also offer a TFBS identification method to scan the presence of identified motifs in their corresponding ChIP-Seq peaks. The major difference is that M was replaced by all peaks in that particular ChIP-Seq dataset and *m* = |*M*|. The background data *H* was generated by the dinucleotide-preserving shuffle strategy. Another difference is that *f*(*x*) was approximated by a Poisson distribution which has been proved in our previous work [13].

### Direct and indirect TF binding in K562 cell line

Among the 690 ChIP-Seq datasets, 150 of them were generated within the K562 cell line, covering 100 TFs. For these 100 ChIP-ed TFs, their canonical motifs were first collected from literature and publicly available databases [67, 68] (Supplementary Table S1). A total of 100 datasets corresponding to the 100 TFs within the K562 cell line were selected from https://genome.ucsc.edu/ENCODE/downloads.html, based on the “first appear, first selected” strategy. All datasets used in this section refer to these 100 datasets.

The peaks in each dataset were then split into the direct-binding (***D***) and the indirect-binding (***I***) peaks, where ***D*** represents the peaks containing the canonical motif of the corresponding ChIP-ed TF, and ***I*** represents peaks that do not have its canonical motif. The average ratio of ***I*** among the 100 datasets is 72% (**Fig. 4**).

A pair of peaks is so-called co-binding peaks if they have at least one nucleotide overlap in our study. For each pair of TFs, e.g., *tf*_1_ and *tf*_2_, we defined that *tf*_1_ tether to *tf*_2_ if and only if

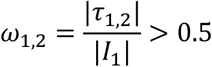

where *I*_1_ indicates the indirect binding peaks of *tf*_1_, τ_1,2_ indicates the co-binding peaks of *I*_1_ and *D*_2_, and *D*_2_ represents the direct binding peaks of *tf*_2_. A total of 61 tethering binding interactions were observed, and 45 of them are MAX-related interactions (**Fig. 4**).

### Shape motif prediction

The prediction of shape motifs followed similar same strategy as above, except that *M* and *H* should be replaced by the corresponding DNA shape features. The mean of shape motif instances was used to generate the shape motif logo, which is represented by a curve. To highlight the difference between shape motif and its background context, its flanking regions (50 bps) were also showcased. A sequence motif logo of underlying sequences of shape motif instances was shown at the lower-left corner of shape motif logo.

### Applied GCNN model to TF-DNA binding specificity prediction

Similar to the CNN model construction above, each input DNA sequence was converted to a *n* × 4 matrix *M* in one-hot format, where *n* indicates the length of the input sequence. The *M* was then fed into the convolutional layer with multiple convolutional filters *E*_*k*_ for *k* = 1,…,16, where each kernel is an *l* × 4 weight matrix. The activation score of each filter on *M* is given by,

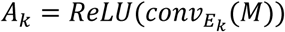

After that, *A*_*k*_ was downsampled by the max pooling layer which has pooling window size *h* × 1 and step size *h* = 10, giving rise to *P*_*k*_ ∈ ℝ^*d*×1^, where 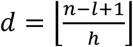. The *P* was reshaped to a *d* × *k* × 1 matrix indicated by *X*, which was then fed into the hidden layers *H*_*i*_ for *i* = 0,…,*L*, where w= 3. The *H*_*i*_ was obtained by:

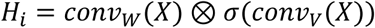

where *W* and *V* are convolutional filters, *σ* a indicates the sigmoid function, and ⊗ is used to calculate the element-wise product. The first two hidden layers have a bottleneck architecture [98] and the output from the last hidden layer was downsampled by a max pooling layer, giving rise to φ. The φ was then fed into two fully connected layers:

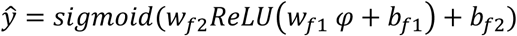

where *w*_*f*1_ and *w*_*f*2_ represent the weights, while *b*_*f*1_ and *b*_*f*2_ indicate the bias units.

## ACKNOWLEDGEMENTS

The authors would like to thank Cankun Wang, Anjun Ma, Shaopeng Gu, Shihan Wu, and Minxuan Sun, for their assistance in pipeline testing and web server development. Especially, the authors thank Prof. Quan Zou from Department of Computer Science at Tianjin University for his insightful discussions and valuable comments on the deep learning model.

## AUTHOR CONTRIBUTIONS

Q.M. and B.Q. conceived the basic idea and designed the overall analyses. J.Y. and Q.M. designed specific experiments, developed the DL framework and carried out most of the computational analysis and data interpretation. J.Y., A.D.H., B.L., and Q.M. wrote the manuscript.

